# Characteristics predicting reduced penetrance variants in the high-risk cancer predisposition gene *TP53*

**DOI:** 10.1101/2025.05.06.652567

**Authors:** Cristina Fortuno, Marcy E. Richardson, Tina Pesaran, Kelly McGoldrick, Paul A. James, Amanda B. Spurdle

**Affiliations:** Population Health Program, QIMR Berghofer, Herston, QLD, 4006, Australia; Ambry Genetics, Aliso Viejo, California, USA; Parkville Familial Cancer Centre, Peter MacCallum Cancer Centre and Royal Melbourne Hospital, Melbourne, VIC, Australia; Sir Peter MacCallum Department of Oncology, University of Melbourne, Melbourne, VIC, Australia; Faculty of Medicine, The University of Queensland, Brisbane, QLD, 4006, Australia

## Abstract

Disease-causing variants with penetrance that is lower than the average expected for a given gene complicate classification, even when using gene-specific guidelines. For *TP53*, a gene associated with some of the highest cancer risks, even reduced penetrance disease-predisposition variants remain clinically actionable. We conducted a review of ClinVar submissions to identify *TP53* variants flagged as having reduced penetrance by genetic testing laboratories and analyzed functional, bioinformatic, frequency, and clinical features of these variants compared to standard pathogenic and benign variants. Our findings show that reported reduced penetrance *TP53* variants are more likely to exhibit intermediate functional activity in multiple assays and are predicted as deleterious with bioinformatic tools, though with lower scores than pathogenic variants. These variants also have a higher population frequency than pathogenic variants, and heterozygotes tend to manifest disease later in life, suggesting a need for refined clinical criteria to better capture attenuated Li-Fraumeni syndrome phenotypes. Finally, by applying a random forest prediction model to all *TP53* uncertain or conflicting variants in ClinVar, we identified additional variants with potential reduced penetrance.

## Introduction

The American College of Medical Genetics and Genomics/Association for Molecular Pathology (ACMG/AMP) germline variant classification guidelines are inherently dichotomous, distinguishing between benign and pathogenic variants for genes associated with Mendelian diseases.^1^ For hereditary cancer predisposition genes associated with high risk of cancer, such as *BRCA1, BRCA2, CDH1, PTEN*, and *TP53*, Clinical Genome (ClinGen) Variant Curation Expert Panels (VCEPs) have helped refine these general guidelines by incorporating gene-specific considerations,^2,3,4,5^ such as establishing expectations based on the standard penetrance associated with a gene of interest.

Disease-causing variants with atypical penetrance, particularly those with reduced penetrance compared to the standard penetrance expected for that gene, pose challenges in variant classification, including when following gene-specific VCEP guidelines. These challenges may include incorrect assumptions about clinical presentation, conflicting or intermediate functional results, and unusual variant frequencies in cases and controls. As a result, these variants are at risk of being classified as variants of uncertain significance (VUS) due to conflicting evidence or, perhaps misclassified as Benign/Likely benign due to the strong weight placed on evidence types that lack sensitivity to distinguish them as reduced penetrance, such as functional data and observation in unaffected individuals. Misclassification as Benign/Likely benign is particularly concerning in clinical settings, as these variants are less likely to be revisited or further investigated compared to VUS simply because of resource constraints. The problem is so significant that ClinGen has set up a Low Penetrance/Risk Allele working group to establish a standardized framework and terminology for categorizing risk alleles and low-penetrance variants in any disease gene (https://www.clinicalgenome.org/site/assets/files/4531/clingenrisk_terminology_recomendations-final-02_18_20.pdf), and the Cancer Variant Interpretation Group-UK has developed a classification framework for cancer risk genes specifically.^6^ While these efforts mark significant progress in the field, there is still a need for more practical evidence-based guidance.

A good example for investigating variants associated with reduced penetrance is the gene *TP53* (HGNC:11998), the cause of Li-Fraumeni syndrome (LFS) (MONDO:0018875), known to have a very strong association with high cancer risks at early ages.^7^ *TP53* germline pathogenic variants are usually linked to LFS core cancers, including early-onset breast cancer, brain cancer, adrenocortical carcinoma, and sarcomas.^8^ This expected clinical presentation, in addition to abundant functional data,^9,10,11,12^ and the existence of *TP53* Variant Classification Expert Panel (VCEP) specifications,^5^ make *TP53* an ideal candidate for investigating features that can better predict variants associated with reduced or atypical cancer penetrance.

Multigene panel testing is increasing the identification of *TP53* germline variants in patients who do not meet the traditional clinical criteria for testing i.e. Classic LFS and Chompret 2015.^8,13^ This may reflect that the spectrum of LFS cancers, and/or their age at presentation, is broader than previously recognized.^14,15^ However, it is also possible that the proportion of *TP53* variants exhibiting reduced penetrance (compared to the average penetrance expected from historical studies) is higher than is currently appreciated. At this time, the only firmly established reduced penetrance *TP53* variant is NM_000546.6(TP53):c.1010G>A (p.R337H), a Brazilian founder variant.^16^ Although several other variants have been proposed as suspected reduced penetrance based on clinical presentation and/or functional results, their age-specific cancer risks remain unconfirmed. Accurate identification of reduced penetrance *TP53* variants, accompanied by age-specific risks, is critical to ensure appropriate patient care.^17^

The aim of this work was to assess the functional, bioinformatic, frequency, and clinical characteristics of a subset of *TP53* pathogenic germline variants reported to exhibit reduced penetrance, to identify attributes that can distinguish them from both *TP53* pathogenic variants with standard penetrance, and from well-established benign variants that are not clinically actionable. We demonstrate that this approach can yield information useful to build future variant classification models capable of separating pathogenic variants with standard penetrance from those associated with reduced penetrance.

## Materials and Methods

### Definitions

In this study, we used the following terminology to categorize variants: benign variants - unlikely to be clinically actionable for *TP53*-related diseases using current specifications; reduced penetrance pathogenic variants - disease-causing but known or suspected to exhibit lower, moderate, or atypical penetrance compared to standard pathogenic *TP53* variants; and pathogenic variants - assumed to have standard penetrance for *TP53*.

### ClinVar review

To identify potential reduced penetrance *TP53* variants, in addition to information available in the literature, we downloaded ClinVar^18^ submissions for *TP53* variants (as at March 14, 2025), and identified evidence summaries that included the terms “reduced”, “moderate”, or “lower penetrance/risk”. At that time, and even at this date (May 7, 2025), there are no ClinVar submissions for *TP53* variants formally categorized as “Pathogenic, low penetrance” or “Established risk allele”. Variants identified in this step formed our reduced penetrance variant group.

### Statistical analyses

#### Distribution of functional results, bioinformatic predictions and allele frequency for pre-defined variant sets

We defined reference sets of *TP53* variants based on ClinVar classifications (as at April 24, 2024), based on VCEP or multiple non-conflicting submissions, as follows: benign reference set, 62 unique missense variants classified as Benign/Likely benign; pathogenic reference set, 113 unique missense variants classified as Pathogenic/Likely pathogenic. Reduced penetrance variants were excluded from these reference sets, and grouped separately regardless of their ClinVar classification.

We then investigated the distribution of functional, bioinformatic, and frequency data for missense variants across the three groups of variants (benign and pathogenic reference sets, and the reduced penetrance variant group as defined above). For functional scores, we used data from currently available systematic assays used by the *TP53* VCEP (specifications v2.3.0), namely Kato et al., 2003,^9^ Giacomelli et al., 2018,^10^ Kotler et al., 2018,^11^ and Funk et al., 2025.^12^ Score cutoffs used to assign loss-of-function (LOF) or noLOF were as defined by the original studies, and currently used by the *TP53* VCEP. We also analysed the Kato data using published data that converted the results into four classes with different levels of functional

disruption (A-D), using a hierarchical Ward’s clustering method.^19^ For bioinformatic scores, we used predictions from BayesDel^20^ and aGVGD^21,22^ which are currently used by the *TP53* VCEP,^5^ and for which we applied the VCEP-defined cutoffs for predicting deleteriousness (where relevant), as well as the more recent predictor AlphaMissense^23^, using the cutoffs defined in the original study. Additionally, we investigated the distribution of the total allele frequency across all genetic ancestry groups for all variants using gnomAD v4.1.^24^

We conducted Kruskal-Wallis tests with post-hoc pairwise comparisons using Wilcoxon tests, and chi- square tests to analyse statistical differences between all variant groups in RStudio (R version 4.4.1).

#### Personal cancer history analysis of clinical presentation for pathogenic versus reduced penetrance variants

We used previously established methods^25^ to conduct personal history analyses using binary logistic regression in SPSS (version 23). Proband data from Ambry Genetics was derived from a multigene panel testing dataset, details of which have been published previously.^26^ After excluding individuals identified to have a *TP53* variant with variant allele fraction <35% (to remove potential mosaic or somatic cases), the Ambry dataset included 256,868 probands negative for Pathogenic/Likely pathogenic or uncertain variants in all genes included in multigene panel tests (as classified by Ambry Genetics), 203 probands with *TP53* reduced penetrance variants (as described above), and 505 probands with *TP53* Pathogenic/Likely pathogenic variants (as classified by Ambry Genetics, excluding those falling in the reduced penetrance group). Of the 189 unique variants observed in the pathogenic group, 59 overlapped with the ClinVar pathogenic reference set.

For this study we conducted two independent analyses, comparing clinical presentation in the baseline group of individuals without a known *TP53* pathogenic variant to: (i) features of individuals with a pathogenic variant; (ii) features of individuals with a reduced penetrance variant.

The phenotypic predictors included in the analysis were the LFS core cancers across different age ranges at first diagnosis, as follows: breast cancer <31y, breast cancer 31-35y, breast cancer 36-60y, brain tumor ≤45y, brain tumor 46-60y, sarcoma ≤45y, sarcoma 46-60y, and adrenocortical carcinoma ≤60y.

Significant differences in the proportion of individuals affected, and their age at first cancer diagnosis, among the three groups were analyzed with chi-squared and Kruskal-Wallis tests, respectively.

#### Random forest predictive model

We built a random forest classification model to categorize the missense variant class (pathogenic, benign, or reduced penetrance) based on scores from the four different functional

assays (Kato, Giacomelli, Kotler, Funk), and the three computational tools (BayesDel, AlphaMissense, and aGVGD) analysed in this study, as well as the total allele frequency. The model was trained using data from the ClinVar reference set variants as well as the reduced penetrance group (excluding c.1101-1G>A), where the missing values in the Kotler and Funk scores were imputed by replacing them with the median value within each variant class. We used 500 trees in the random forest, and model performance was evaluated using a confusion matrix. The feature importance of each score was also assessed to understand which variables contributed most to the model predictions using Mean Decrease in Gini values.

We applied this model to an additional group of 12 Benign/Likely benign and 21 Pathogenic/Likely pathogenic missense variants in ClinVar classified by single submitters only (and therefore not included in the original ClinVar reference sets), as well as 497 *TP53* missense variants classified as Uncertain significance or with conflicting classifications in ClinVar (as at March 14, 2025), in order to identify new potential reduced penetrance *TP53* variants based on their unique functional, bioinformatic and allele frequency characteristics. All of these 530 variants had values for all components in the model.

## Results

### Suspected reduced penetrance variants

In addition to the known Brazilian founder NM_000546.6(TP53):c.1010G>A (p.R337H) variant,^16^ we identified 10 more variants (9 missense, 1 splice site) listed in ClinVar meeting our search criteria for designation as (suspected) reduced penetrance (Table 1). An additional variant, NM_000546.6(TP53):c.467G>A (p.R156H), was included due to being flagged as suspected reduced penetrance by Couch et al., 2017^27^ and Behoray et al., 2022.^28^ Only four of the 11 variants were classified as Pathogenic/Likely pathogenic in ClinVar, one of these being the Brazilian founder variant. Rationale for inclusion of these variants, including ClinVar evidence summaries and/or literature is detailed in Supplementary Table 1.

**Table 1.**
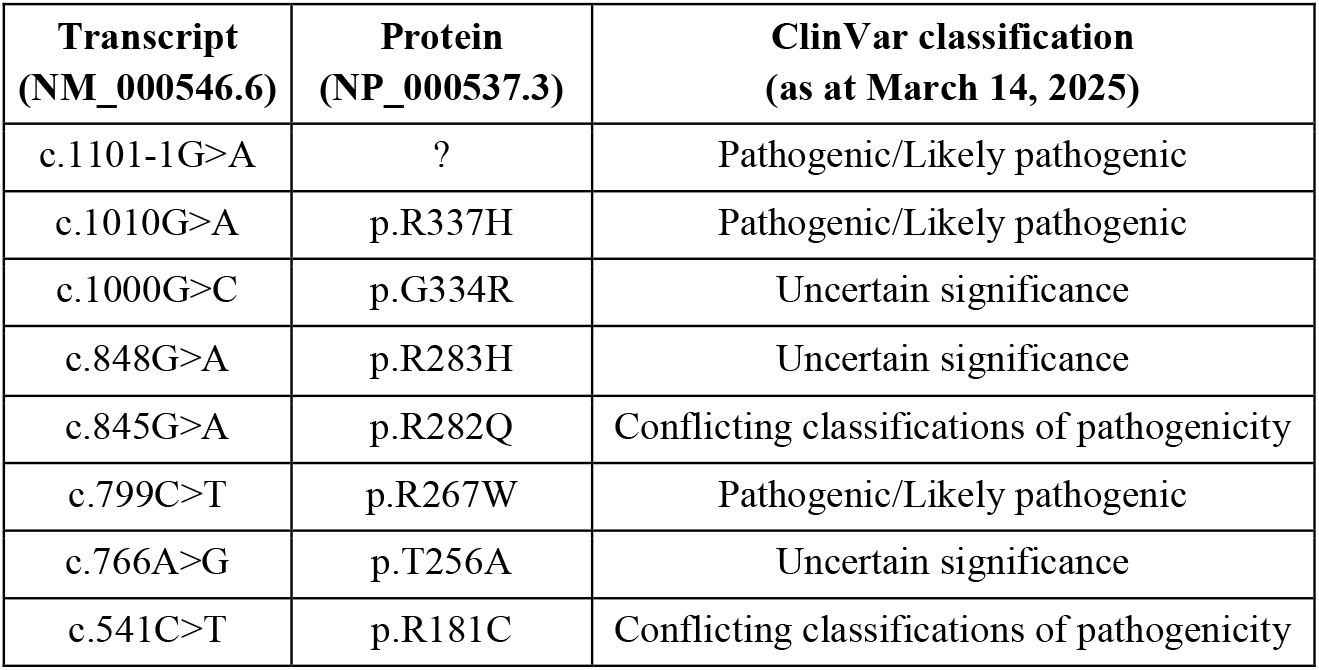
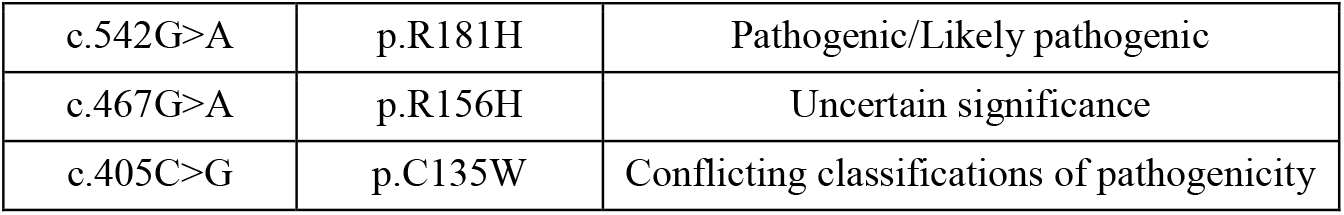
*TP53* variants included in the reduced penetrance group.

### Comparison of functional impact for missense variants

Kato and Giacomelli assay results were available for all 185 missense variants across the three variant groups. Kotler assay data was available for 137 variants (including eight with reduced penetrance), and Funk assay data for 134 variants (including eight with reduced penetrance), due to both of these assays being restricted to specific exons. The vast majority (>85%) of variants in the benign or pathogenic reference sets exhibited functional impact corresponding to noLOF or LOF, respectively (Table 2). Of the four functional assays, only Kato defined a score range indicating intermediate function; seven out of 10 (70%) reduced penetrance missense variants fell within this range, with LOF category assigned to two of the three remaining variants in this group. For the remaining assays, LOF was assigned for one (Giacomelli), two (Kotler), and six (Funk) of the reduced penetrance variants.

**Table 2.** Median functional scores for all variant groups and number of variants in each functional category per variant group.

| Variant group |  | Kato | Giacomelli | Kotler | Funk |
| --- | --- | --- | --- | --- | --- |
| Total variants assessed |  | 185 | 185 | 137 | 134 |
| Median functional score<br>(n noLOF, n partial function, n LOF) | Benign | 93.70<br>(49, 12, 1) | 0.54<br>(61, na, 1) | -2.09<br>(20, na, 0) | -0.86<br>(18, na, 1) |
|  | Reduced | 40.30<br>(1, 7, 2) | 0.19<br>(9, na, 1) | -1.82<br>(6, na, 2) | 0.66<br>(2, na, 6) |
|  | Pathogenic | 6.60<br>(0, 4, 109) | -1.00<br>(9, na, 104) | 0.05<br>(5, na, 104) | 0.93<br>(1, na, 106) |
| P-value* | | $1.67 \times 10^{-28}$ | $7.20 \times 10^{-29}$ | $4.51 \times 10^{-13}$ | $3.11 \times 10^{-12}$ |
| Cutoff for noLOF |  | >75 | >-0.21 | ≤-1 | <0 |
| Partial function range |  | 20-75 | na | na | na |
| Cutoff for LOF |  | ≤20 | ≤-0.21 | >-1 | ≥0 |

LOF = Loss-of-function
\*Kruskal-Wallis test comparing differences across the three groups of variants

Differences in the distribution of functional assay scores across all three variant groups were highly significant (p <0.001) (Table 2). Overall, the median distribution of functional assay scores for the reduced penetrance variants fell between that of benign and pathogenic variants (Figure 1). The distribution of Kato and Giacomelli scores differed highly significantly between pairwise comparisons of the variant groups (p <0.001). Regarding the Kotler scores, significant differences were observed between the benign vs. pathogenic and reduced penetrance vs. pathogenic groups (p <0.001), but no significant difference was found between the benign vs. reduced penetrance groups (p >0.05). For the Funk assay, differences in scores were highly significant comparing benign vs. pathogenic or reduced penetrance (p <0.001), with somewhat smaller differences between the reduced penetrance vs. pathogenic group (p <0.01).

**Figure 1.**
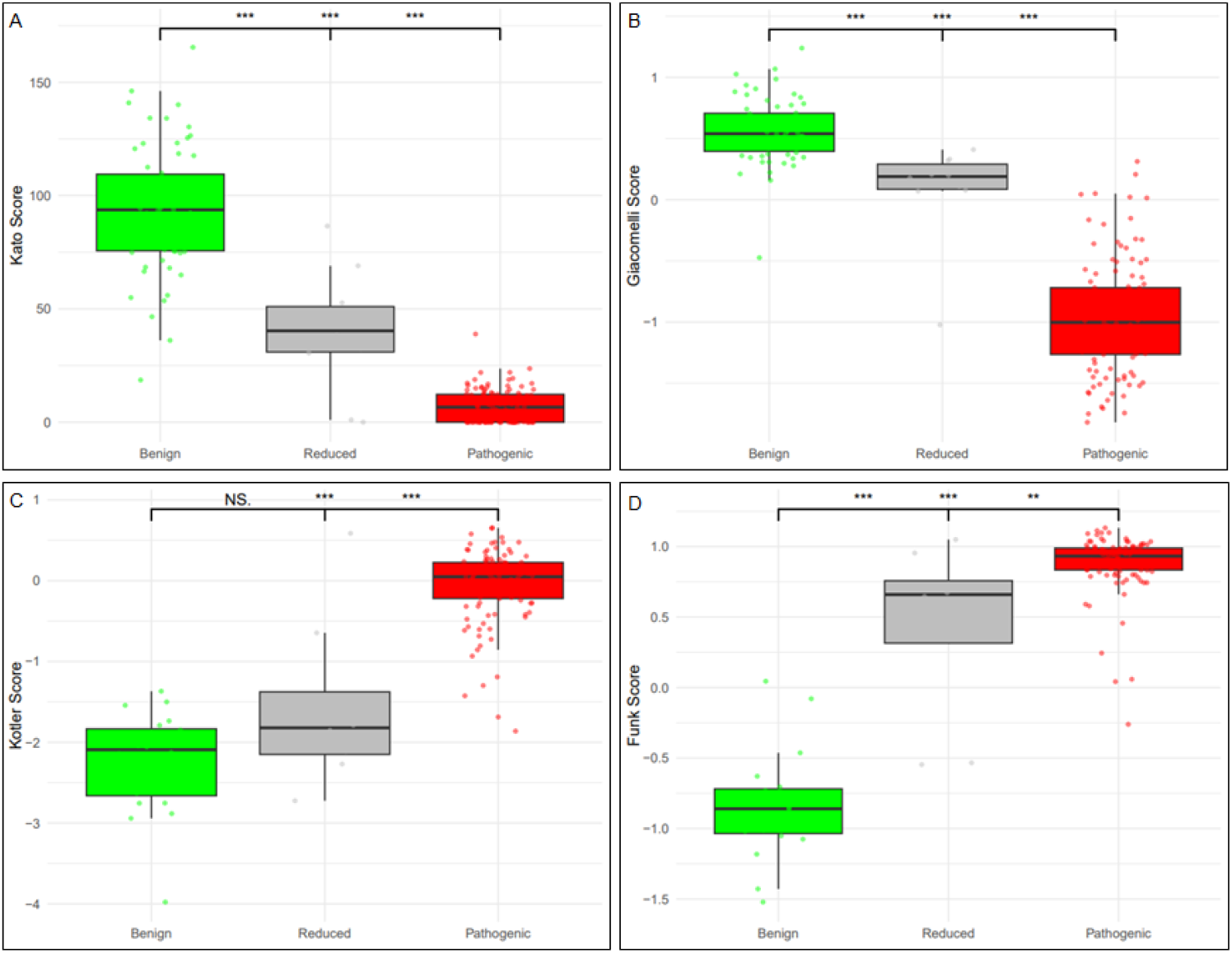
Distribution of functional assay scores across *TP53* benign (green), reduced penetrance (grey), and pathogenic (red) variants. A: Kato et al., 2003, B: Giacomelli et al., 2018, C: Kotler et al., 2018, D: Funk et al., 2025. P-values refer to pairwise comparisons with Wilcoxon tests, where * = p < 0.05, ** = p < 0.01, and *** = p < 0.001.

Statistical comparisons noted in the center are for Benign versus Pathogenic variants Significant differences in the distribution of the Kato clustering classes were observed across the three variant groups (p = 1.41×10^−35^). The benign group was predominantly represented by class D (42/62, 68%), the reduced penetrance group by class C (7/10, 70%), and the pathogenic group by class A (93/113, 82%) (Figure 2).

**Figure 2.**
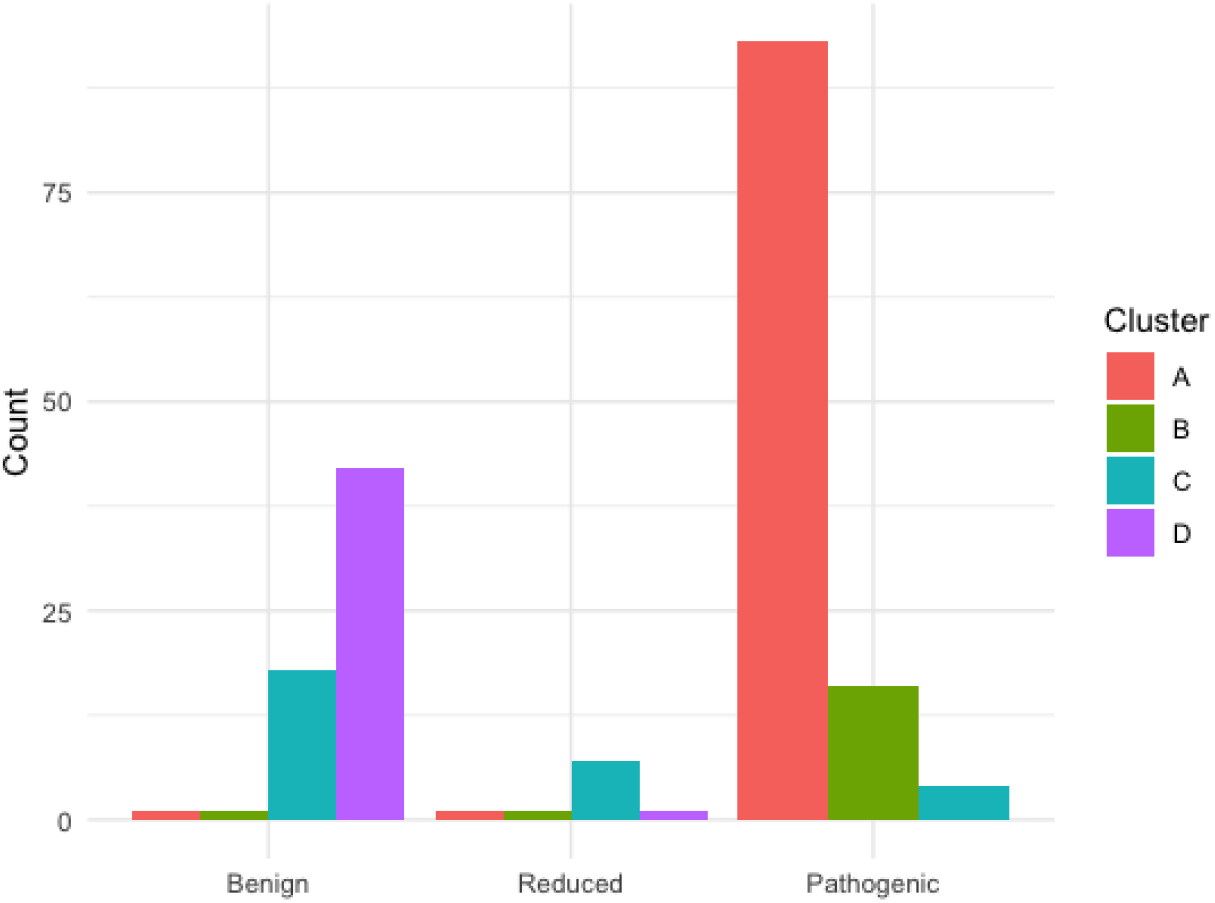
Distribution of Kato clustering classes across *TP53* benign, reduced penetrance, and pathogenic variants. (p = 1.41e-35). Classes represent a gradient of yeast-based transcriptional activity, from lowest (A) to highest (D) as per Montellier et al., 2024

### Bioinformatic analysis of missense variants

Bioinformatic predictions were available for all 185 missense variants in all three variant groups. Both the BayesDel and AlphaMissense tools performed well in predicting deleteriousness for benign and pathogenic variants, with median scores typically below or above the selected cutoffs, respectively (Table 3). All variants in the reduced penetrance and pathogenic groups were predicted as deleterious by BayesDel, while 54 of the 62 (87%) benign variants were predicted neutral. For AlphaMissense, which includes an ambiguous category, five out of 62 (8%) benign variants, seven out of 10 (70%) reduced penetrance variants, and 110 out of 113 (97%) pathogenic variants were predicted deleterious. Two variants in each group were predicted to have an intermediate effect. Only one variant in both the reduced penetrance and pathogenic groups was predicted as neutral, compared to 55 (89%) in the benign group.

**Table 3.** Median bioinformatic scores for all variant groups and number of variants in each bioinformatic category per variant group.

| Variant group |  | BayesDel | AlphaMissense |
| --- | --- | --- | --- |
| Total variants assessed |  | 185 | 185 |
| Median bioinformatic score (n predicted neutral, n intermediate, n deleterious) | Benign | -0.05<br>(54, na, 8) | 0.08<br>(55, 2, 5) |
|  | Reduced | 0.47<br>(0, na, 10) | 0.82<br>(1, 2, 7) |
|  | Pathogenic | 0.55<br>(0, na, 113) | 0.99<br>(1, 2, 110) |
| P-value* | | $2.45 \times 10^{-27}$ | $1.58 \times 10^{-27}$ |
| Cutoff for neutral |  | <0.16 | <0.34 |
| Intermediate range |  | na | 0.564-0.34 |
| Cutoff for deleterious | | $\geq 0.16$ | >0.564 |

\*Kruskal-Wallis test comparing differences across the three groups of variants

Differences in the distribution of bioinformatic results were highly significant across all three variant groups (p <0.001) (Table 3). Similar to observations for the functional assays, the median distribution of BayesDel and AlphaMissense scores for reduced penetrance variants appeared to be positioned between that of benign and pathogenic variants, and distributions differed highly significantly across all three variant groups (p <0.001) (Figure 3). For both tools, pairwise comparisons showed that the differences between reduced penetrance and pathogenic variants were not as marked (p <0.05 for BayesDel and p <0.01 for AlphaMissense) as for the remaining comparisons (p <0.001).

**Figure 3.**
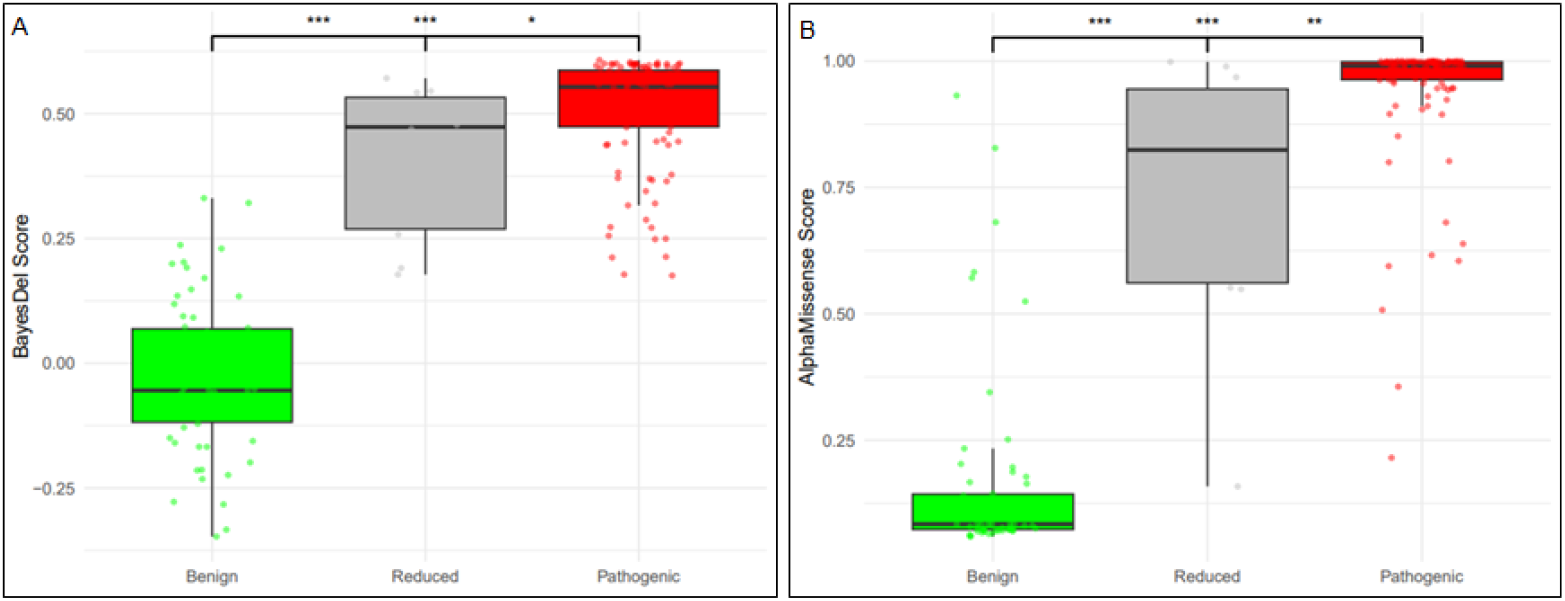
Distribution of bioinformatic scores across *TP53* benign (green), reduced penetrance (grey), and pathogenic (red) variants. **A: BayesDel** (Feng et al., 2017), **B: AlphaMissense** (Cheng et al., 2023). P-values refer to pairwise comparisons with Wilcoxon tests, where * = p < 0.05, ** = p < 0.01, and *** = p < 0.001. Statistical comparisons noted in the center are for Benign versus Pathogenic variants

The distribution of the aGVGD categories was significantly different across the three variant groups (p = 5.31×10^−22^). The benign group was predominantly represented by class C0 (54/62, 87%), the pathogenic group by class C65 (65/113, 57%), while the reduced penetrance group had a higher diversity of categories (Figure 4). In particular, three reduced penetrance variants were C0 (predicted neutral), three were C65 (predicted most deleterious), while the other four fell into categories between.

**Figure 4.**
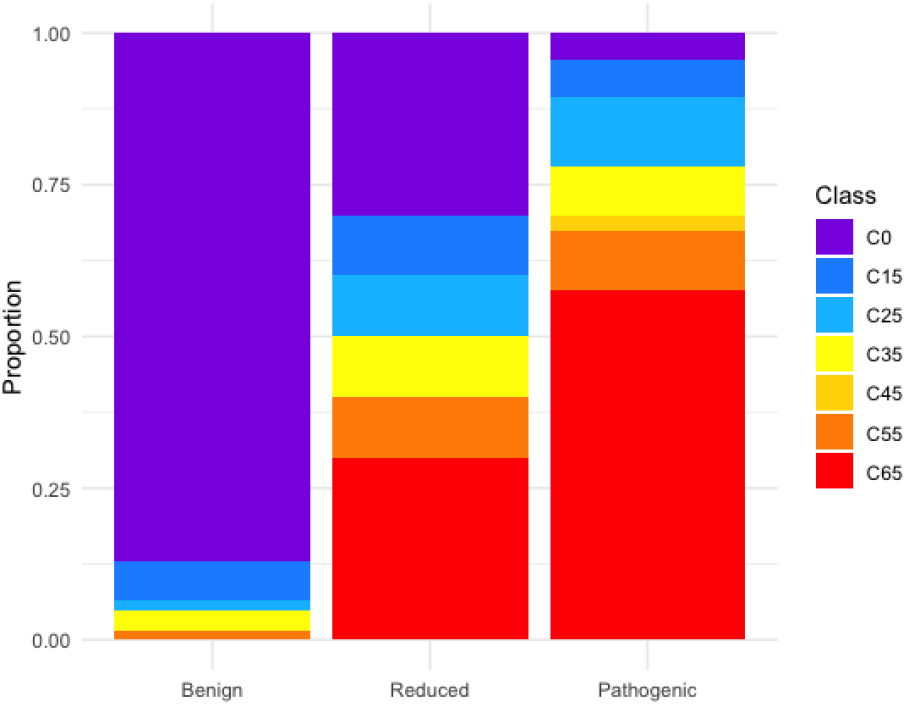
Distribution of aGVGD categories across *TP53* benign, reduced penetrance, and pathogenic variants. (p = 5.31×10^−22^). C65 is predicted most likely to interfere with function, and C0 least likely, based on the biochemical variation at each position in a multiple sequence alignment of orthologous sequences as per Tavtigian et al., 2008

### Differences in allele frequency across groups

The distribution of allele frequency differed significantly across all three variant groups (p = 1.126×10^−05^) (Table 4). All 113 pathogenic variants met PM2_Supporting (as defined by the *TP53* VCEP specifications v2.3.0), as opposed to nine of 11 (82%) reduced penetrance variants and 40 of 62 (64%) benign variants. Of note, six of the variants in the benign group met BS1 based on the filtering allele frequency (as defined by the *TP53* VCEP specifications v2.3.0), as opposed to none in the reduced penetrance groups. As expected, benign variants exhibited the highest allele frequency, pathogenic variants the lowest, and reduced penetrance variants fell in between (Figure 5). The differences between benign and pathogenic variants, as well as between reduced penetrance and pathogenic variants, were both highly significant (p < 0.001). However, the allele frequency distribution between benign and reduced penetrance variants did not differ significantly (p > 0.05).

**Table 4.** Total allele frequency from gnomAD v4.1 for all variant groups (n=186 total variants) and number of variants meeting population frequency codes per variant group.

| Variant group | Allele frequency median | N meeting PM2_Supporting | N meeting any benign code (BA1 or BS1) |
| --- | --- | --- | --- |
| Benign | 0.0000068 | 40 | 6 (BS1) |
| Reduced | 0.0000019 | 9 | 0 |
| Pathogenic | 0 | 113 | 0 |
| P-value* | $1.13 \times 10^{-05}$ | | |
| Cutoff for PM2_Supporting | Total allele frequency $< 0.00003$ (and $< 0.00004$ for any single genetic ancestry group with multiple alleles in non-founder populations) | | |
| Cutoff for BS1 | Filtering allele frequency in non-founder populations $\geq 0.0003$ | | |

\*Kruskal-Wallis test comparing differences across the three groups of variants

**Figure 5.**
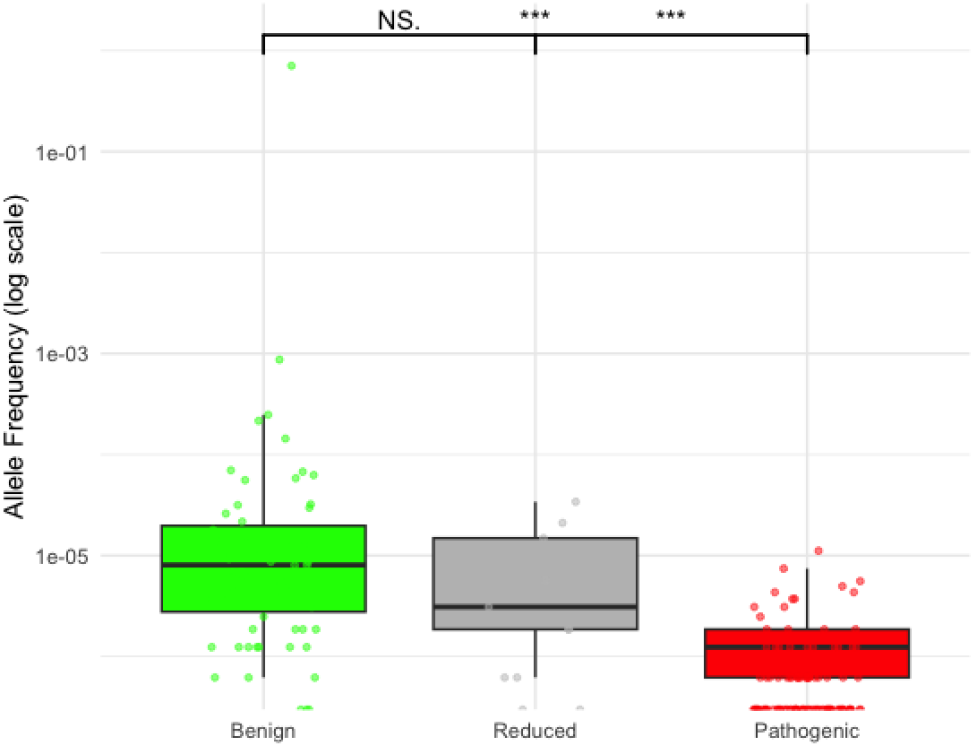
Distribution of total allele frequency in gnomAD v4.1 across *TP53* benign (green), reduced penetrance (grey), and pathogenic (red) variant groups (p = 1.13×10^05^)

### Phenotypic predictors associated with reduced penetrance variants

The Ambry dataset used for the personal history analysis had proband data for carriers of all reduced penetrance variants included in this study. The variants with the highest number of probands were c.542G>A (p.R181H) and c.467G>A (p.R156H), with 46 probands each, while c.405C>G (p.C135W) and c.1101-1G>A had only one proband each. The number of individuals with the remaining reduced penetrance variants ranged from two to 28. For this analysis, the control group included individuals without a known Pathogenic/Likely pathogenic or uncertain variant.

The proportion of individuals affected with any cancer for the control, reduced penetrance, and pathogenic groups was 40%, 43%, and 48%, respectively (p <0.001), with average age at first cancer diagnosis being 50.3y, 47.7y, and 36.6y, respectively (p <0.001).

In the phenotypic prediction analysis, ORs represent enrichment of the phenotype within the cohort of tested individuals, and not cancer risk relative to baseline population risk. The ORs indicated significant enrichment of the following phenotypes in the pathogenic variant group relative to individuals without a pathogenic variant in this cohort (Table 5): breast cancer <31y, breast cancer 31-35y, brain tumor ≤45y, sarcoma ≤45y, sarcoma 46-60y, and adrenocortical carcinoma ≤60y. In the reduced penetrance group, only breast cancer <31y and sarcoma ≤45y showed significant enrichment compared to individuals without a pathogenic variant, but with much lower ORs compared to the pathogenic variant group (OR 3.05 vs. 16.07 for breast cancer <31y, significantly different based on the confidence intervals; OR 7.09 vs. 27.85 for sarcoma ≤45y, not significantly different). The reduced penetrance group lacked enough data for sarcoma 46-60y and adrenocortical carcinoma ≤60y.

**Table 5.** ORs associated with core LFS cancers among *TP53* pathogenic and reduced penetrance variants against 256,868 controls*.

| Phenotype (1<sup>st</sup> cancer) | Pathogenic group (n=505) |  |  |  |  | Reduced penetrance group (n=203) |  |  |  |
| --- | --- | --- | --- | --- | --- | --- | --- | --- | --- |
|  | P-value | OR | Lower 95% CI | Upper 95% CI |  | P-value | OR | Lower 95% CI | Upper 95% CI |
| **Breast cancer <31y** | <0.001 | 16.07 | 12.08 | 21.38 |  | 0.01 | 3.05 | 1.25 | 7.45 |
| **Breast cancer 31-35y** | <0.001 | 3.57 | 2.36 | 5.42 |  | 0.95 | 0.97 | 0.31 | 3.03 |
| **Breast cancer 36-60y** | 0.41 | 0.90 | 0.70 | 1.16 |  | 0.89 | 1.03 | 0.72 | 1.47 |
| **Brain tumor ≤45y** | <0.001 | 14.40 | 6.36 | 32.60 |  | 0.10 | 5.19 | 0.72 | 37.27 |
| **Brain tumor 46-60y** | No data |  |  |  |  | No data |  |  |  |
| **Sarcoma ≤45y** | <0.001 | 27.85 | 16.92 | 45.85 |  | <0.01 | 7.09 | 1.75 | 28.73 |
| **Sarcoma 46-60y** | <0.001 | 20.34 | 8.29 | 49.92 |  | No data |  |  |  |
| **Adrenocortical carcinoma ≤60y** | 0.01 | 12.29 | 1.69 | 89.22 |  | No data |  |  |  |

CI = Confidence interval; OR = Odds ratio (enrichment of the phenotype within these case-case cohorts, and not cancer risk relative to baseline population risk)
\*Controls include probands with no Pathogenic/Likely pathogenic or uncertain variants identified via multigene panel testing

### Identification of additional potential reduced penetrance *TP53* missense variants

The random forest model was trained to categorize variants into three classes (pathogenic, benign, and reduced penetrance), based on eight components (Kato, Giacomelli, Kotler, Funk, BayesDel, AlphaMissense, aGVGD, and total allele frequency). The model performed well in predicting benign (1.6% classification error) and pathogenic variants (0.9% classification error), but showed a higher error rate for reduced penetrance variants (30%), as shown by the confusion matrix (Supplementary Table 2).

The feature importance analysis of the random forest model revealed that the Kato score was the most influential feature in categorizing variants into the three groups, with the highest Mean Decrease in Gini value of 19.9, closely followed by Funk (19.6) (Supplementary Figure 1). Other influential features were Kotler (17.2) and Giacomelli (16.8). Allele frequency had the lowest importance in this model, with a Mean Decrease in Gini value of 1.4.

**Supplementary Figure 1.**
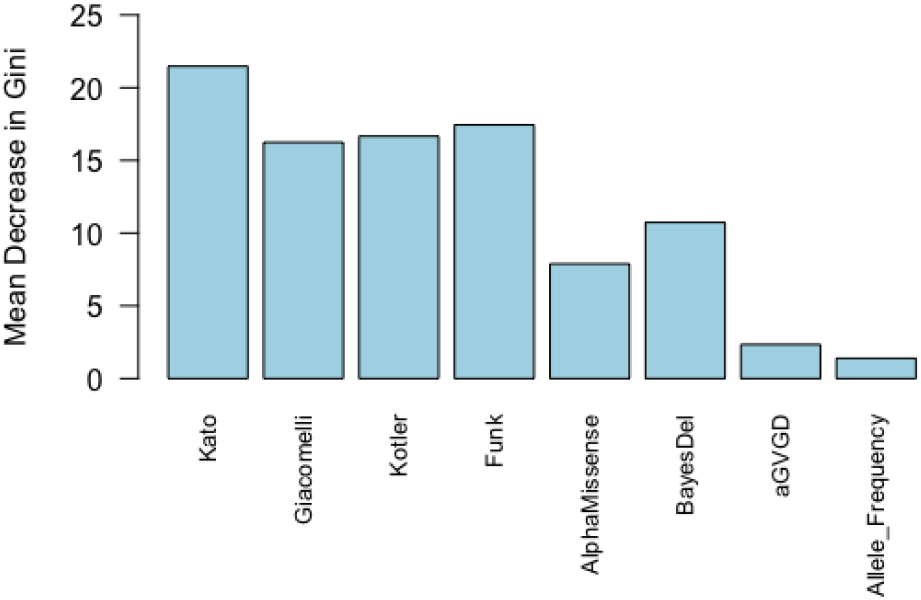
Mean Decrease in Gini values representing the importance for the different components included in the random forest model to predict pathogenic, benign, and reduced penetrance variant status

When this model was applied to additional variants in ClinVar (that is, variants not in any of the reference groups), the model categorization agreed with classifications for all 12 Benign/Likely benign variants and all 21 Pathogenic/Likely pathogenic single submissions in ClinVar. Of the remaining 497 Uncertain significance or conflicting *TP53* missense variants to which this model was applied, 60 were categorized as reduced penetrance, of which 51 were Uncertain and nine conflicting in ClinVar (Supplementary Table 3).

## Discussion

For most cancer genes, current variant classification models are not designed to classify reduced penetrance variants. For genes like *TP53*, where pathogenic variants are linked to an extremely high cancer risk,^7^ and have associated intensive surveillance strategies,^29^ even variants with a lower-than-average penetrance warrant clinical action. Further, for variants that are already classified as pathogenic, it is important to confirm which of these have reduced penetrance, as this could lead to different clinical management strategies for carriers, as it is the case for R337H carriers.^30^

Our study enhances understanding of the functional, bioinformatic, frequency, and clinical characteristics of reported reduced penetrance cancer gene variants, using *TP53* as an example, and demonstrates how these features can be used to provide a foundation for prediction models to better identify reduced penetrance disease-causing variants as being distinct from benign or standard penetrance pathogenic variants. Importantly, based on the distribution of these features, we developed a preliminary predictive model that identified 60 potential reduced penetrance *TP53* variants, currently in ClinVar as VUS or conflicting. We provide these variants in Supplementary Table 3 as examples of those that could be prioritized for further analyses using future *TP53*-specific reduced penetrance models and/or formal penetrance studies. While we highlight the preliminary nature of this predictive model, based on a small reference set of reported (suspected) reduced penetrance variants, we note that the model correctly identified all variants with single submission assertions as Benign/Likely benign or Pathogenic/Likely pathogenic.

From examination of the different data types, we highlight several interesting observations which collectively suggest that existing ClinGen VCEP approaches could be extended to better identify reduced penetrance variants, by incorporating more precise functional ranges, recalibrated bioinformatic tools, and descriptions of attenuated LFS phenotypes that can better distinguish between reduced and standard penetrance variants.

First, reduced penetrance variants were more likely to exhibit intermediate functional activity, as demonstrated by four independent functional studies.^9,10,11,12^ Among existing assays, only Kato explicitly includes a “partial function” category, where most reduced penetrance variants clustered in this study. While this could have served as a potential reason for them being flagged as potential reduced penetrance by the ClinVar submitters, similar distributions were also observed for the other functional assays. Notably, the Kato clustering classes,^19^ offering greater granularity than the original three Kato categories, could be leveraged for improved specificity, in line with a recent cancer risk study which found that carriers of cluster B or C variants were less likely to meet clinical testing criteria for LFS.^31^ Ideally, new functional assays should be developed to better capture intermediate functions or alternative disease mechanisms, but our findings suggest that existing assays could be recalibrated for finer granularity to identify suspected reduced penetrance *TP53* variants. Further, current assays differ slightly in how they analyze functional impact, and our study revealed distinct differences in the distribution of results across the three variant groups, indicating that integrating multiple assays into a combined model may be more informative than relying on a single assay, as has been suggested in the context of multiplexed assays of variant effect (Calhoun et al., submitted).^32^

Second, reduced penetrance variants tended to be predicted as deleterious by multiple bioinformatics tools, although with lower scores than pathogenic variants on average. In addition, aGVGD^21,22^ displayed a greater diversity of results for reduced penetrance variants compared to the other groups. Although AlphaMissense^23^ currently includes a score range for intermediate function, most reduced penetrance variants were predicted deleterious following the cutoffs recommended in the original publication, suggesting that recalibration of this tool to introduce more categories could enhance its value to separate reduced penetrance from standard penetrance pathogenic variants. Similarly, recalibration to define additional categories for BayesDel^20^ could also improve its utility for identifying variants with reduced penetrance.

Third, the total allele frequency distribution of reduced penetrance variants more closely resembled that of benign variants than pathogenic variants, although most still met the PM2_Supporting criterion, suggesting that their somewhat higher frequency would not impact the application of pathogenic clinical codes (i.e., PS4) which require PM2_Supporting to be met in order to be considered.

Last, as we would expect for reduced penetrance variants, in comparison to standard pathogenic variants we observed later average age at first cancer diagnosis, and the core LFS cancers currently used as clinical evidence of pathogenicity were not as highly associated. However, it is notable that the most characteristic *TP53*-associated cancers, particularly early-onset breast cancer and sarcomas, remained significantly associated with reduced penetrance variants, although with lower level of enrichment than for the group with standard pathogenic variants. This all suggests that more attenuated LFS phenotypes should be considered for application of clinical evidence in favor of pathogenicity to facilitate identification of *TP53* reduced penetrance variants.^14^

We recognize the potential for circularity in our study. Our reference sets of pathogenic and benign variants were derived from ClinVar, where some of the predictive components analyzed in this study would have contributed to their classifications. However, no single component alone would have been decisive in classifications made by Expert Panels or through multiple non-conflicting submissions, as these typically rely on a combination of evidence types. Additionally, we incorporated newer functional assays^12^ and bioinformatic tools^23^ that were published after our reference sets were established, ensuring that these classifications were not directly influenced by these newer methodologies. In addition, as previously noted, some components used in this study, such as Kato results, may have contributed to flagging certain variants as potentially reduced penetrance. However, the similar distribution observed across other functional studies suggests that this is not solely driven by the Kato results. Suspicions of reduced penetrance are typically supported by internal clinical data from laboratories, rather than relying on functional results alone, as indicated by the evidence summaries in ClinVar (Supplementary Table 1). Additionally, the fact that our model predictions aligned correctly with all single pathogenic and benign submissions in ClinVar is not unexpected, given that components of the model likely influenced the original classification of these variants. Nevertheless, this performance demonstrates the model’s ability to align with these classifications and to prioritize other potential reduced penetrance variants for research purposes.

Another substantial limitation is the small sample size for the reduced penetrance group, particularly given that most of these variants are suspected, with only one being officially confirmed.^16^ It is possible that some of these variants may not truly exhibit reduced penetrance. It is also conceivable that some variants included in the pathogenic and benign reference sets could be associated with reduced penetrance. In the end, both the low number of reduced penetrance variants and uncertainty in the reference panels reflect the difficulty of identifying these variants with certainty, an issue that our model aims to begin to address.

Future larger studies with a greater number of variants will be essential to validate and confirm our findings. Additional larger studies will also play a key role in quantifying age-specific risks for reduced penetrance variants highlighted in this study, aiming to improve the clinical management of carriers and enhance the accuracy of cancer risk prediction for affected individuals and their relatives.

## References

1. Richards, S., Aziz, N., Bale, S., Bick, D., Das, S., Gastier-Foster, J., Grody, W.W., Hegde, M., Lyon, E., Spector, E., et al. (2015). Standards and guidelines for the interpretation of sequence variants: a joint consensus recommendation of the American College of Medical Genetics and Genomics and the Association for Molecular Pathology. Genetics in medicine : official journal of the American College of Medical Genetics 17, 405–424.

2. Parsons, M.T., de la Hoya, M., Richardson, M.E., Tudini, E., Anderson, M., Berkofsky-Fessler, W., Caputo, S.M., Chan, R.C., Cline, M.S., Feng, B.J., et al. (2024). Evidence-based recommendations for gene-specific ACMG/AMP variant classification from the ClinGen ENIGMA BRCA1 and BRCA2 Variant Curation Expert Panel. Am J Hum Genet 111, 2044–2058.

3. Luo, X., Maciaszek, J.L., Thompson, B.A., Leong, H.S., Dixon, K., Sousa, S., Anderson, M., Roberts, M.E., Lee, K., Spurdle, A.B., et al. (2023). Optimising clinical care through CDH1-specific germline variant curation: improvement of clinical assertions and updated curation guidelines. Journal of medical genetics 60, 568–575.

4. Mester, J.L., Ghosh, R., Pesaran, T., Huether, R., Karam, R., Hruska, K.S., Costa, H.A., Lachlan, K., Ngeow, J., Barnholtz-Sloan, J., et al. (2018). Gene-specific criteria for PTEN variant curation: Recommendations from the ClinGen PTEN Expert Panel. Human mutation 39, 1581–1592.

5. Fortuno, C., Lee, K., Olivier, M., Pesaran, T., Mai, P.L., de Andrade, K.C., Attardi, L.D., Crowley, S., Evans, D.G., Feng, B.J., et al. (2021). Specifications of the ACMG/AMP variant interpretation guidelines for germline TP53 variants. Human mutation 42, 223–236.

6. Garrett, A., Allen, S., Durkie, M., Burghel, G.J., Robinson, R., Callaway, A., Field, J., Frugtniet, B., Palmer-Smith, S., Grant, J., et al. (2025). Classification of variants of reduced penetrance in high-penetrance cancer susceptibility genes: Framework for genetics clinicians and clinical scientists by CanVIG-UK (Cancer Variant Interpretation Group-UK). Genetics in medicine : official journal of the American College of Medical Genetics 27, 101305.

7. Fortuno, C., Feng, B.J., Carroll, C., Innella, G., Kohlmann, W., Lázaro, C., Brunet, J., Feliubadaló, L., Iglesias, S., Menéndez, M., et al. (2024). Cancer Risks Associated With TP53 Pathogenic Variants: Maximum Likelihood Analysis of Extended Pedigrees for Diagnosis of First Cancers Beyond the Li-Fraumeni Syndrome Spectrum. JCO Precis Oncol 8, e2300453.

8. Li, F.P., and Fraumeni, J.F., Jr. (1969). Soft-tissue sarcomas, breast cancer, and other neoplasms. A familial syndrome? Ann Intern Med 71, 747–752.

9. Kato, S., Han, S.Y., Liu, W., Otsuka, K., Shibata, H., Kanamaru, R., and Ishioka, C. (2003). Understanding the function-structure and function-mutation relationships of p53 tumor suppressor protein by high-resolution missense mutation analysis. Proceedings of the National Academy of Sciences of the United States of America 100, 8424–8429.

10. Giacomelli, A.O., Yang, X., Lintner, R.E., McFarland, J.M., Duby, M., Kim, J., Howard, T.P., Takeda, D.Y., Ly, S.H., Kim, E., et al. (2018). Mutational processes shape the landscape of TP53 mutations in human cancer. Nature genetics 50, 1381–1387.

11. Kotler, E., Shani, O., Goldfeld, G., Lotan-Pompan, M., Tarcic, O., Gershoni, A., Hopf, T.A., Marks, D.S., Oren, M., and Segal, E. (2018). A Systematic p53 Mutation Library Links Differential Functional Impact to Cancer Mutation Pattern and Evolutionary Conservation. Mol Cell 71, 178–190 e178.

12. Funk, J.S., Klimovich, M., Drangenstein, D., Pielhoop, O., Hunold, P., Borowek, A., Noeparast, M., Pavlakis, E., Neumann, M., Balourdas, D.I., et al. (2025). Deep CRISPR mutagenesis characterizes the functional diversity of TP53 mutations. Nature genetics 57, 140–153.

13. Bougeard, G., Renaux-Petel, M., Flaman, J.M., Charbonnier, C., Fermey, P., Belotti, M., Gauthier-Villars, M., Stoppa-Lyonnet, D., Consolino, E., Brugieres, L., et al. (2015). Revisiting Li-Fraumeni Syndrome From TP53 Mutation Carriers. Journal of clinical oncology : official journal of the American Society of Clinical Oncology 33, 2345–2352.

14. Kratz, C.P., Freycon, C., Maxwell, K.N., Nichols, K.E., Schiffman, J.D., Evans, D.G., Achatz, M.I., Savage, S.A., Weitzel, J.N., Garber, J.E., et al. (2021). Analysis of the Li-Fraumeni Spectrum Based on an International Germline TP53 Variant Data Set: An International Agency for Research on Cancer TP53 Database Analysis. JAMA Oncol 7, 1800–1805.

15. de Andrade, K.C., Khincha, P.P., Hatton, J.N., Frone, M.N., Wegman-Ostrosky, T., Mai, P.L., Best, A.F., and Savage, S.A. (2021). Cancer incidence, patterns, and genotype-phenotype associations in individuals with pathogenic or likely pathogenic germline TP53 variants: an observational cohort study. The Lancet Oncology 22, 1787–1798.

16. Pinto, E.M., and Zambetti, G.P. (2020). What 20 years of research has taught us about the TP53 p.R337H mutation. Cancer 126, 4678–4686.

17. Moghadasi, S., Meeks, H.D., Vreeswijk, M.P., Janssen, L.A., Borg, A., Ehrencrona, H., Paulsson-Karlsson, Y., Wappenschmidt, B., Engel, C., Gehrig, A., et al. (2018). The BRCA1 c. 5096G>A p.Arg1699Gln (R1699Q) intermediate risk variant: breast and ovarian cancer risk estimation and recommendations for clinical management from the ENIGMA consortium. Journal of medical genetics 55, 15–20.

18. Landrum, M.J., Lee, J.M., Riley, G.R., Jang, W., Rubinstein, W.S., Church, D.M., and Maglott, D.R. (2014). ClinVar: public archive of relationships among sequence variation and human phenotype. Nucleic Acids Res 42, D980–985.

19. Montellier, E., Lemonnier, N., Penkert, J., Freycon, C., Blanchet, S., Amadou, A., Chuffart, F., Fischer, N.W., Achatz, M.I., Levine, A.J., et al. (2024). Clustering of TP53 variants into functional classes correlates with cancer risk and identifies different phenotypes of Li-Fraumeni syndrome. iScience 27, 111296.

20. Feng, B.J. (2017). PERCH: A Unified Framework for Disease Gene Prioritization. Human mutation 38, 243–251.

21. Tavtigian, S.V., Byrnes, G.B., Goldgar, D.E., and Thomas, A. (2008). Classification of rare missense substitutions, using risk surfaces, with genetic- and molecular-epidemiology applications. Human mutation 29, 1342–1354.

22. Fortuno, C., James, P.A., Young, E.L., Feng, B., Olivier, M., Pesaran, T., Tavtigian, S.V., and Spurdle, A.B. (2018). Improved, ACMG-Compliant, in silico prediction of pathogenicity for missense substitutions encoded by TP53 variants. Human mutation.

23. Cheng, J., Novati, G., Pan, J., Bycroft, C., Žemgulytė, A., Applebaum, T., Pritzel, A., Wong, L.H., Zielinski, M., Sargeant, T., et al. (2023). Accurate proteome-wide missense variant effect prediction with AlphaMissense. Science 381, eadg7492.

24. Chen, S., Francioli, L.C., Goodrich, J.K., Collins, R.L., Kanai, M., Wang, Q., Alföldi, J., Watts, N.A., Vittal, C., Gauthier, L.D., et al. (2024). A genomic mutational constraint map using variation in 76,156 human genomes. Nature 625, 92–100.

25. Fortuno, C., Michailidou, K., Parsons, M., Dolinsky, J.S., Pesaran, T., Yussuf, A., Mester, J.L., Hruska, K.S., Hiraki, S., O’Connor, R., et al. (2024). Challenges and approaches to calibrating patient phenotype as evidence for cancer gene variant classification under ACMG/AMP guidelines. Human molecular genetics 33, 724–732.

26. Fortuno, C., McGoldrick, K., Pesaran, T., Dolinsky, J., Hoang, L., Weitzel, J.N., Beshay, V., San Leong, H., James, P.A., and Spurdle, A.B. (2022). Suspected clonal hematopoiesis as a natural functional assay of TP53 germline variant pathogenicity. Genetics in medicine : official journal of the American College of Medical Genetics 24, 673–680.

27. Couch, F.J., Shimelis, H., Hu, C., Hart, S.N., Polley, E.C., Na, J., Hallberg, E., Moore, R., Thomas, A., Lilyquist, J., et al. (2017). Associations Between Cancer Predisposition Testing Panel Genes and Breast Cancer. JAMA Oncol 3, 1190–1196.

28. Nehoray, B., Schwartz, A., Hyman, S., Stokes, S., Cervantes, A., Amos, C., Gruber, S.B., and Garber, J.E. (2022). Questioning a Li-Fraumeni Syndrome diagnosis: Characterization of a commonly observed germline <i>TP53</i> variant, p.Arg156His. Journal of Clinical Oncology 40, 10612–10612.

29. Achatz, M.I., Villani, A., Bertuch, A.A., Bougeard, G., Chang, V.Y., Doria, A.S., Gallinger, B., Godley, L.A., Greer, M.-L.C., Kamihara, J., et al. (2025). Update on Cancer Screening Recommendations for Individuals with Li–Fraumeni Syndrome. Clinical Cancer Research, OF1–OF10.

30. Galante, P.A.F., Guardia, G.D.A., Pisani, J., Sandoval, R.L., Barros-Filho, M.C., Gifoni, A., Patrão, D.F.C., Ashton-Prolla, P., de Vasconcellos, V.F., Freycon, C., et al. (2025). Personalized screening strategies for TP53 R337H carriers: a retrospective cohort study of tumor spectrum in Li-Fraumeni syndrome adult carriers. Lancet Reg Health Am 42, 100982.

31. Müntnich, L.J., Dutzmann, C.M., Großhennig, A., Härter, V., Keymling, M., Mastronuzzi, A., Montellier, E., Nees, J., Palmaers, N.E., Penkert, J., et al. (2025). Cancer risk in carriers of TP53 germline variants grouped into different functional categories. JNCI Cancer Spectr 9.

32. Calhoun, J.D., Dawood, M., Rowlands, C.F., Fayer, S., Radford, E.J., McEwen, A.E., Turnbull, C., Spurdle A.B., Starita L.M., Jagannathan S. (2025). Combining multiplexed functional data to improve variant classification. arXiv.

